# Structural basis for the ubiquitin chain recognition of the human 26S proteasome

**DOI:** 10.1101/2025.04.07.647569

**Authors:** Sascha J. Amann, Robert Kalis, Maximilian Fottner, Hannah Knaudt, Irina Grishkovskaya, Johannes Zuber, Kathrin Lang, Nicholas G. Brown, David Haselbach

## Abstract

Proteasomal degradation is a fundamental process for all eukaryotic life. A protein destined for degradation is first tagged with a polyubiquitin chain, which is selected by the proteasome. Different ubiquitin chain topologies serve as distinct signals, with K48-linked chains acting as the canonical degradation signal and K11/K48-branched chains providing even more potent targeting, particularly during cell cycle regulation. However, the structural basis for how the proteasome distinguishes between these different chain architectures has remained unclear. Here, we present high-resolution cryo-EM structures of the human 26S proteasome bound to both a K48-linked tetraubiquitin chain and a K11/K48-branched chain. Our structures reveal distinct binding modes for these two types of chain linkage. K48 chains wrap around the Ubiquitin interaction motif of the receptor RPN10 in an unexpected spiral conformation, while K11 branches engage the proteasome through previously uncharacterised interfaces in a cleft formed between RPN2 and RPN10. Through structure-guided mutagenesis and cellular studies, we demonstrate that these binding modes are essential for efficient substrate degradation and cell cycle progression. These findings establish how the proteasome achieves selective substrate recognition through chain topology-specific interactions.

---

The ubiquitin chain architecture dictates the fate of more than 80% of cellular proteins through the ubiquitin proteasome system (UPS). Theoretically out of four ubiquitin moieties more than 1000 different chain topologies could arise through distinct lysine residues (K6, K11, K27, K29, K33, K48, K63) or the N-terminal methionine (M1). Amongst these numerous possible topologies of ubiquitin chains, only a subset with specific linkage types and length leads to efficient degradation. K48-linked tetraubiquitin chains represent the canonical degradation signal (*1, 2*), while K11/K48-branched chains serve as potent accelerators of degradation during cell cycle progression (*3, 4*) as they are the predominant chain type generated by the cell cycle regulator APC/C. Eventually, the human 26S proteasome, the cell’s primary protein degradation machinery, must distinguish between an array of ubiquitin chain topologies to ensure precise proteostasis.(*5*) However, the structural principles underlying the 26S proteasome’s selectivity for specific ubiquitin chain topologies remain largely unknown.

The 26S proteasome consists of the proteolytically active 20S core particle (CP) and the 19S regulatory particle (19S), which recognizes and unfolds ubiquitinated substrates.(*6*) A typical proteasome substrate contains both a unstructured region and a ubiquitin chain (*7, 8*). The unstructured region is required to engage with the proteasomes ATPase and initialize unfolding of the target protein. The ubiquitin chain is required for substrate recognition. Currently, three well-characterized ubiquitin receptors have been identified within the 19S regulatory particle: the proteasomal subunits RPN1 (*9*), RPN10 (*10*), and RPN13 (*11*), each containing distinct ubiquitin-binding domains that recognize individual ubiquitins. Additionally, the intrinsic deubiquitinating enzyme RPN11 can bind ubiquitin and may act as a molecular switch controlling proteasome function (*12*). How these individual binding sites contribute to the recognition of specific ubiquitin chains, however, remained unclear. Intriguingly, a systematic mutation study in *Saccharomyces cerevisiae* revealed that even when these main receptors are deleted, significant ubiquitin recognition capability remained, suggesting the existence of additional, yet unidentified receptors (*13*).

Several structural studies on proteasomes have revealed many aspects of substrate degradation (*14*–*17*). However, only the ubiquitin moiety proximal to the substrate could be resolved interacting with the deubiquitinase RPN11, leaving unclear how different ubiquitin chain topologies are recognized by the proteasomal ubiquitin receptors. Here, we present high-resolution cryo-EM structures of the 26S proteasome bound to both a homotypic K48-linked tetra-ubiquitin chain and a K11/K48-branched ubiquitin chain, providing unprecedented insights into how different ubiquitin chain topologies are recognized and degraded by the proteasome. These structures reveal the molecular basis for chain topology-specific recognition and suggest mechanisms for how the proteasome integrates multiple binding interactions to achieve selective degradation of ubiquitinated substrates.

## Results

### Structure of K48-linked Tetraubiquitin Bound to the Proteasome

Previous structural studies that provided significant insights into proteasomal function (*14, 15*) employed K63-linked chains, which are known to be involved in other cellular processes than proteasomal degradation. However, structures with K48-linked chains, the canonical degradation signal, have remained elusive. Our initial attempts revealed only single ubiquitin moieties rather than complete chains (*18*), highlighting two key challenges: the rapid deubiquitination of K48 chains (occurring within seconds) (*19*) and the heterogeneity of naturally formed chains in both attachment sites and length. To overcome this challenge, we used our Ubl-tool strategy combining enzymatic assembly, genetic code expansion and sortase-mediated transpeptidation to generate a defined homotypically K48-tetraubiquitinated target protein with a non-hydrolysable linkage between a specific lysine in a target protein and the most proximal Ub moiety. (*20, 21*) The multistep assembly process is based on the UBE2R1-mediated generation of a native K48-diUb (diUb-A) and a K48-linked diUb (diUb-B) bearing a Sortase2A motif at the C-terminus of the proximal Ub, which are subsequently joined in a second UBE2R1-mediated reaction to access a K48-linked tetraUb bearing a Sortase2A recognition motif at the proximal Ub (Fig. 1a, fig. S1a). By using Sortase2A, we charged the K48-linked tetraUb onto an sfGFP bearing the non-canonical amino acid GGisoK at position 215 and a CycB degron at its C-terminus as unstructured proteasome initiation site (Fig. S1a, fig. S1b). This resulted in a defined homotypically K48-linked tetraUb sfGFP-K215GGisoK-CycB conjugate, that is recalcitrant towards cleavage by deubquitinases due to the two point mutations (R72A and R74T) introduced at the C-terminus of the most proximal Ub to allow for Sortase2A recognition.

**Fig. 1.**
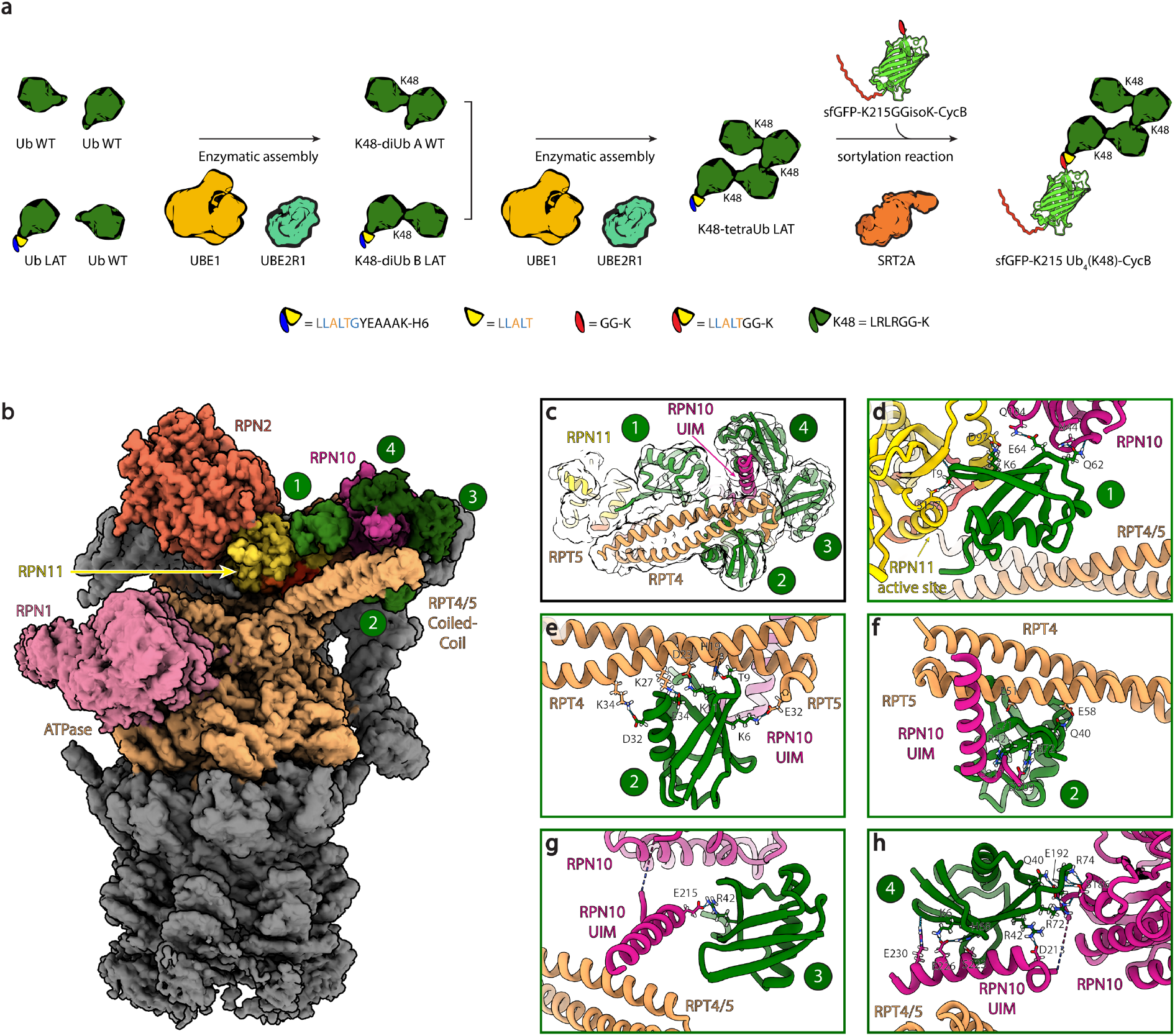
Human 26S proteasome interacting with a homotypic K48-linked ubiquitin chain. (a) Generation of a defined homotypic K48-tetraUb-GFP conjugate. Enzymatically generated diUbs-A/B are combined for the assembly of a homotypic K48-tetraUb bearing a Sortase 2A motif at the C-terminus of the most proximal Ub. K48-tetraUb is charged via sortase-mediated transpeptidation onto the non-canonical amino acid bearing sfGFP-K215GGK-CycB resulting in homotypic K48-tetraUb-sfGFP-K215GGK-CycB conjugate used for CryoEM studies. (b) CryoEM density of human 26S proteasome interacting with a homotypic K48 linked tetra-ubiquitin chain (1-4) with colored subunits. (c) K48-linked tetra ubiquitin chain model fitted in gauss filtered CryoEM density. (d-h) Models of the proximal (d), second (e-f), third (g), and fourth (h) ubiquitins of the K48 linked ubiquitin chain interacting with the 26S proteasome

Using this biochemically homogeneous K48-linked tetraubiquitin chain conjugate, we determined a single-particle cryo-EM structure of the ubiquitin chain bound to the 26S proteasome at a resolution of 2.9 Å (Fig. 1b, fig. S2, Table S1). The proteasome is in a translocation incompetent pre-engagement state termed S1. We observed distinct densities for all four ubiquitin moieties, with resolution gradually decreasing from the substrate proximal to distal ubiquitin moieties (Fig. 1c, fig S2d). While the three proximal ubiquitins (Ub1, Ub2, Ub3) could be modeled unambiguously into their corresponding densities, the most distal ubiquitin (Ub4) was positioned based on its previously reported interaction with the ubiquitin-interacting motif 1 (UIM1) of RPN10, the canonical ubiquitin receptor (*22*) (pdb 1yx5).

Unexpectedly, the K48-linked tetraubiquitin chain adopts a spiral conformation as it wraps around the RPN10^UIM1^ helix (Fig 1b, c). The most proximal ubiquitin (Ub1) interacts with the metalloprotease domain of the proteasomal deubiquitinase RPN11 (Fig. 1d). This Ub1 carries two engineered point mutations (R72A, R74T), which prevent its full engagement with RPN11 and subsequent deubiquitination, likely enabling the structural determination of the entire K48-linked tetra-ubiquitin chain (Fig. 1c). However the comparison with previously determined fully engaged RPN11–ubiquitin complexes (*14, 15*) suggests that the observed chain configuration represents a physiologically relevant pre-engagement state and provides the structural basis for the initial proteasomal recognition of K48-linked ubiquitin chains. (Supplementary Movie S1). Due to the missing ability of deubiquitinate we could resolve now also in the context of the intact proteasome.

The second most proximal ubiquitin (Ub2) establishes interactions with the coiled-coil domain formed by RPT4 and RPT5 subunits, consistent with previous genetic studies (*23*) suggesting this region as a potential ubiquitin receptor (Fig. 1e, f). Additionally we observed that Ub2 also interacts with RPN1^UIM1^, essentially crosslinking coiled coil and the UIM, explaining why deletion of RPN1^UIM1^ leads to a less resolved Rpt4/5 coiled coil (*24*). The 3^rd^ proximal ubiquitin (Ub3) seems to only loosely interact but again with the RPN10^UIM1^ (Fig. 1g).

Finally the most distal ubiquitin (Ub4) also engages with RPN10^UIM1^ but via its canonical binding interface (Fig. 1h) (*22*). Interestingly, three ubiquitins of the chain interact with the RPN10^UIM1^ explaining its importance. Notably, arginine 42 (R42) of ubiquitins 2-4 mediates previously uncharacterized interactions with RPN10^UIM1^ through a network of hydrogen bonds and salt bridges (Fig. 1f, g, h).

### Structure of the K11/K48 branched chain on the proteasome

In addition to K48-linked tetra-ubiquitin chains, recent studies have identified branched K11/K48-linked ubiquitin chains as particularly effective degradation signals. (*3, 25*) These branched chains are primarily synthesized by the Anaphase Promoting Complex/Cyclosome (APC/C) and are essential for cell cycle progression.

To generate a substrate bearing a K11/K48-branched ubiquitin chain for structural analysis, we used a recombinant system comprising the APC/C and its E2 enzymes, UBE2C and UBE2S, to polyubiquitinate the native substrate Securin. UBE2C initiates ubiquitination by priming the substrate with K48-linked chains, while UBE2S extends these chains by conjugating K11-linked ubiquitin chains, resulting in branched K11/K48 chains (*26*–*31*) (Fig. 2a, fig. S1c, d).

**Fig. 2.**
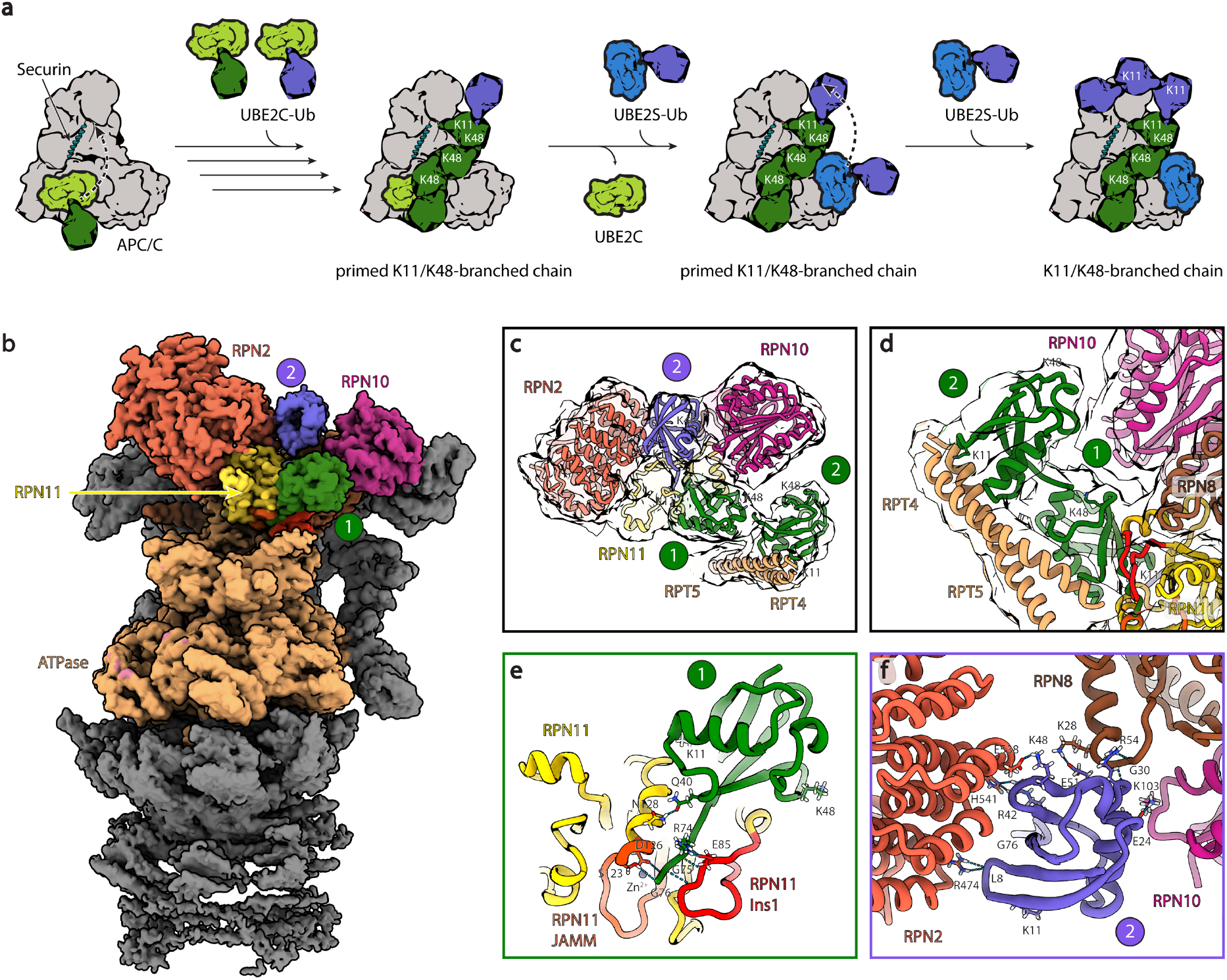
Human 26S proteasome interacting with a heterotypic K11/K48 branched ubiquitin chain. (a) Scheme of how APC/C forms K11/K48 branched chains on Securin. (b) CryoEM density of human 26S proteasome interacting with a K11 linked ubiquitin chain (1-2) with colored subunits. (c-d) K11/K48 branched chain model fitted in gauss filtered CryoEM density. (e-f) Models of the proximal (d), and second (e) ubiquitin of the K11 linked di-ubiquitin interacting with the 26S proteasome

To gain insights into the mechanistic details of how K11/K48-branched chains engage with the 26S proteasome, we used single-particle cryo-EM and successfully resolved three ubiquitin moieties of the branched chain. (Fig. 2, fig. S3, Table S1). The proximal ubiquitin and the K11-branched ubiquitin were resolved at sufficient high resolution enabling to model side chains densities (Fig. 2b, fig. S3d), while the K48 branch showed lower resolution but still enabled unambiguous fitting (Fig. 2c, d).

The substrate-proximal ubiquitin fully engages with RPN11 similarly to the K48 chain but being competent for deubiquitination (*14, 15*)(Fig 2e). The K11 branch is well-resolved and occupies a specific cleft formed between RPN2 and RPN10 (Fig. 2f). Notably, lysine 11 of the second ubiquitin points away from the proteasome. This observation implies that short mono-ubiquitin branches could be sufficient for recognition. Furthermore, K48 of the K11-linked ubiquitin participates in the interaction with RPN2, suggesting that mixed-linkage chains produced by the APC/C may require an alternative interaction surface.

We also identified previously uncharacterized interactions between the K11-linked ubiquitin with RPN8 and RPN10, revealing a more extensive recognition surface than previously known.

The K48 branch engages the RPT4/5 coiled-coil domain in a manner similar to that observed in the homotypic K48-linked tetraubiquitin chain structure, suggesting additive binding effects for the K11/K48-branched chain. (Fig. 1c, e, f, Fig. 2c, d). The simultaneous interactions of the K11/K48 ubiquitin branches suggest a mechanistic basis for the enhanced recognition of branched chains by the proteasome.

### Genetic validation

To validate our structural findings in a cellular context, we established two complementary cell reporter systems monitoring substrate-specific protein turnover in RKO cells (Fig. 3a). The first system tracks the degradation of the transcription factor c-MYC, which is predominantly targeted through K48-linked ubiquitination. The second system monitors the cell cycle regulator Securin, an APC/C substrate known to be targeted through K11/K48-branched chains. Both proteins were tagged with mCherry to enable quantitative assessment of their steady-state levels. We engineered proteasome variants by overexpressing mutant subunits in wild-type cells, exploiting the endogenous orphan quality control mechanism that ensures quantitative exchange of proteasomal subunits, achieving high incorporation of the variant subunits.

**Fig. 3.**
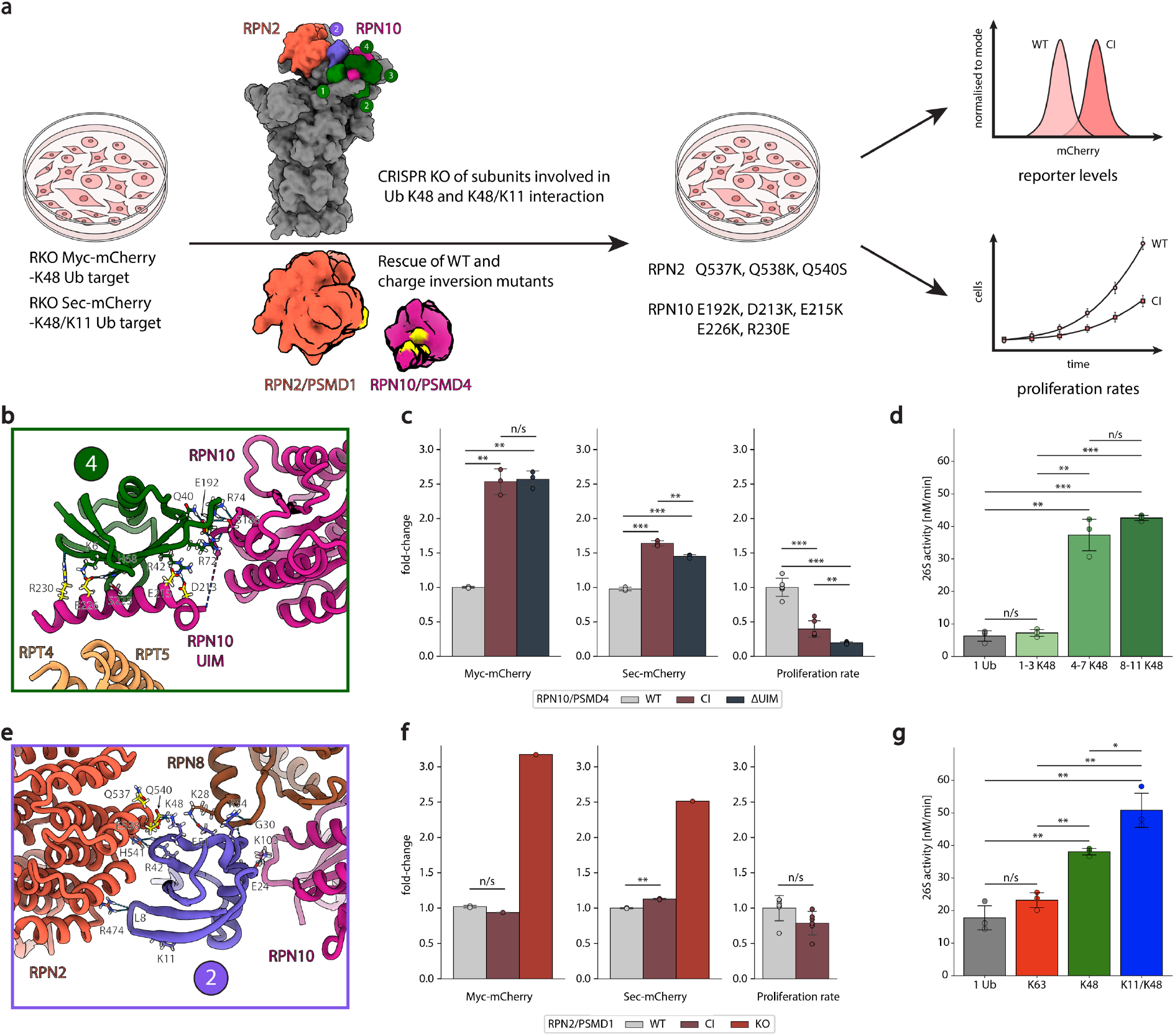
Validation of the ubiquitin chain CryoEM model using functional genetics and kinetic assays. (a) Overview of the functional genetic assay strategy. (b) Model of the distal ubiquitin moiety of a K48-linked tetra ubiquitin chain interacting with the 26S proteasome with indicated mutated residues of RPN10 used for the charge inversion mutant (yellow). (c) Proliferation rates of cells with WT, UIM deletion or charge inversion mutant of PSMD4/RPN10 overexpression and Myc- or Securin-mCherry levels of cell with additional KO of the endogeneous gene (n = 3, error bars indicate ± SD, two-tailed t-test, n.s.: p > 0.05, *: p ≤ 0.05, **: p ≤ 0.01, ***: p ≤ 0.001). (d) 26S proteasomal substrate degradation rates depending on K48-linked ubiquitin chain lengths (n = 3, error bars indicate ± SD, two-tailed t-test, n.s.: p > 0.05, *: p ≤ 0.05, **: p ≤ 0.01, ***: p ≤ 0.001). (e) Model of the distal ubiquitin moiety of a K11-linked di-ubiquitin interacting with the 26S proteasome with indicated mutated residues of RPN2 (yellow) used for the charge inversion mutant in f. (f) Proliferation rates of cells with WT (n = 3), charge inversion mutant (CI, n = 3) or KO only (n = 1) of PSMD1/RPN2 overexpression and Myc- or Securin-mCherry levels of cell with additional KO of the endogeneous gene (error bars indicate ± SD, two-tailed t-test, n.s.: p > 0.05, *: p ≤ 0.05, **: p ≤ 0.01, ***: p ≤ 0.001). (g) 26S proteasomal substrate degradation rates depending on Mono Ub, K63-linked, K48-linked or K11/K48-branched ubiquitin chain (n = 3, error bars indicate ± SD, two-tailed t-test, n.s.: p > 0.05, *: p ≤ 0.05, **: p ≤ 0.01).

To assess the functional significance of the K48-specific binding sites, we introduced mutations in the *PSMD4* gene encoding RPN10, focusing on residues identified in our structural analysis (E192K, D213K, E215K, E226K, R230E) and a *PSMD4* ΔUIM version as positive control (Fig. 3b). When overexpressed in cells, both mutations caused a proliferation defect compared to a wild-type PSMD4 overexpression, indicative of incorporation of the dysfunctional subunit in the proteasome (Fig. 3c left). After knock out of the endogenous *PSMD4* gene, mutant cells showed both elevated MYC and Securin protein levels, consistent with compromised proteasomal degradation (Fig. 3c center, right). Observing this effect upon the disruption of the binding interface of the fourth ubiquitin moiety is in agreement with previous findings that a chain consisting of at least four K48-linked ubiquitin moieties is the minimum requirement for 26S proteasomal degradation (*2*) . Using an in vitro 26S proteasome degradation assay we could also reproduce this (Fig. 3d, fig. S4a, b, fig. S5a, b). Notably, the accumulation of Securin was less pronounced compared to c-MYC, suggesting either differential synthesis rates or the existence of alternative degradation pathways.

To investigate the specific contribution of K11-chain recognition, we mutated the newly identified interaction surfaces in RPN2 (encoded by PSMD1) (Fig. 3e). Compared to an rescued RPN2 WT, these mutations caused no change in Myc levels and led to a modest but significant accumulation of Securin, accompanied by a slight, though non-significant, reduction in proliferation when overexpressed without knockout of the endogenous gene (Fig. 3f). These findings indicate that this single interaction between K48 of the K11-linked ubiquitin and RPN2 is not sufficient to explain the increased levels of substrate degradation of K11/K48 branched chains compared to homotypic K48-linked chains (*4, 25*) . However, using an in vitro 26S proteasome degradation assay we could also reproduce that K11/K48 branched chains are degraded faster than K48-linked chains while both are degraded faster than substrates containing a single ubiquitin moiety or a K63 linked chain (Fig. 3g, fig. S4, fig S4c, d).

## Discussion

### Chain Length Requirements and Dynamic Implications of Ubiquitin Recognition

As recapitulated in this study, efficient proteasomal degradation requires K48-linked ubiquitin chains of at least four ubiquitin moieties. Our structural analysis reveals the molecular basis for this length requirement: only the fourth ubiquitin establishes strong interactions with the RPN10^UIM1^, which serves as the primary recognition adaptor. The functional significance of this interaction is demonstrated by our mutational analysis, where point mutations disrupting the UIM1-distal ubiquitin interface impair c-MYC degradation to the same extent as deletion of the entire C-terminus of RPN10, consisting of the UIM1 and the UIM2 motif. This suggests that RPN10^UIM1^ acts as the pivotal recognition site, with other interactions playing supporting roles. Of note, a recent study indicates that K48-linked chains of only three ubiquitins may be sufficient (*32*). While this contrasts with our data in the cellular context more factors may be involved in the recognition than in our isolated system. For instance, a degradation via a USP14 associated proteasome may use a different recognition mechanism (*16*).

The chain trajectory we observe creates an intriguing link between key regulatory elements of the proteasome. The ubiquitin chain extends from RPN11 through the RPT4/5 coiled-coil to the RPN10^UIM1^, effectively bridging these functionally important regions. This arrangement has potential mechanistic implications for proteasome activation, which requires a substantial ∼30-degree rotation of the regulatory particle lid toward the RPT4/5 coiled-coil, bringing RPN10 and the coiled-coil into closer proximity (*33*). The ubiquitin chain’s tethering of these elements could facilitate this conformational change. While this movement would seemingly require displacement of the chain, the flexible attachment of RPN10^UIM1^ and the RPT4/5 coiled-coil domain provides sufficient conformational freedom to accommodate such rearrangements without disrupting key recognition contacts (Supplementary Movie S2). However, this displacement of RPN10^UIM1^ and the RPT4/5 coiled-coil domain could also represent an additional energy barrier, increasing the substrate’s dwell time prior to the 30° conformational change and thereby preventing premature deubiquitination, which can be observed in the case of K63-linked chains (fig. S5e, f, g).(*32*) The exact trajectory of ubiquitin chain recognition will need to be the subject of future studies.

### From Discrete Receptors to an Integrated Recognition Surface

Previous studies demonstrated that the proteasome retains the ability to recognize polyubiquitinated substrates even when the well-characterized ubiquitin receptors RPN1, RPN10, and RPN13 are mutated out in vitro, suggesting the existence of additional recognition mechanisms. The RPT4/5 coiled-coil domain was previously proposed as an additional receptor (*23*), and our structures confirm interactions with this region.

Our structures reveal a more complex recognition paradigm where ubiquitin chains engage multiple 19S subunits through an extensive network of spatially distributed interactions. Each individual contact contributes relatively few interactions, but collectively they form a larger, composite binding surface that extends beyond discrete receptor subunits. This distributed recognition model is exemplified by RPN10, where we identified multiple points of contact outside its canonical UIM1 domain interaction. The presence of such extensive interaction surfaces may explain the remarkable robustness of ubiquitin recognition by the proteasome, as the loss of individual receptors can be partially compensated by the remaining network of weaker interactions. Importantly, this distributed recognition could serve as a kinetic proofreading mechanism, where multiple weak interactions collectively increase substrate residence time, providing a longer window of opportunity for productive substrate engagement and commitment to degradation.

### Recognition of K11/K48 Branched Chains and Their Regulatory Implications

While homotypic K48 chains constitute the predominant degradation signal in cells, K11/K48 branched chains serve as more potent targeting signals. These branched chains are particularly important for cell cycle regulation, as they are generated by the APC/C(*25*). Our structural analysis reveals that the K48 and K11 branches have distinct binding surfaces, with the K48 branch engaging the same binding sites as homotypic K48 chains. The K11 branch, however, follows a different trajectory, utilizing novel interaction surfaces that were not classified as ubiquitin receptors so far. This additional binding interface effectively expands the total interaction surface area, likely explaining the enhanced binding affinity of branched chains. Interestingly, a parallel study that used a mixed K11/K48 chain finds that a K48 branch on the K11 branch even further binds the upper lid area having even more binding surface available. Indicating tighter binding than with homotypic branches (*34*).

Notably, the trajectory of the K11 branch overlaps with the binding site of PITHD1, a protein we identified as a proteasome inhibitor (*18*). This spatial overlap suggests a mechanism where PITHD1 specifically interferes with the recognition of K11/K48 branched chains, potentially functioning as a mitosis-specific proteasome inhibitor.

Here we provide unprecedented structural insights into the initial recognition of distinct ubiquitin chain topologies by the proteasome, revealing how this cellular machine achieves both specificity and versatility in substrate selection. The distinct binding modes we observed for K48 and K11/K48-branched chains suggest a sophisticated recognition system where different chain architectures engage specific surfaces on the proteasome, likely serving as a critical determinant of degradation efficiency. However, many questions remain about how initial chain recognition is coordinated with subsequent steps of substrate processing. Understanding this progression from recognition to commitment, including the roles of deubiquitinating enzymes, ATPase-driven unfolding, and potential proofreading mechanisms, will be crucial. The existence of chain-specific binding sites also raises the intriguing possibility that other ubiquitin chain topologies may utilize additional, yet undiscovered interaction surfaces. This emerging picture of chain topology-specific recognition not only advances our understanding of cellular protein quality control but also suggests new therapeutic strategies for selectively modulating proteasome function in disease. A new generation of targeted proteolysis inducing drugs may now further take chain topology into account.

## Methods

### Generation of K48-linked tetraubiquitin bearing sfGFP-K215GGK-CycB

Purification of Ub variants, E2 enzymes and Sortase 2A as well as synthesis of AzGGisoK and its site-specific incorporation into amber codon (TAG)-bearing sfGFP (sfGFP-K215TAG-CycB) and its subsequent Staudinger reduction were carried out as previously described (*20, 21*). In short, K48-linked diUb A was generated by incubating 1 mM Ub wt with 5 μM UBE2R1 and 50 nM UBA1 in 50 mM Tris pH 7.5, 10 mM MgCl2-ATP. After overnight incubation at 37 °C the reaction mixture was diluted 1/20 in 50 mM Ammonium acetate pH 4.5 followed by cation exchange using a MonoS column (Cytiva) and size exclusion chromatography using a Superdex 75 16/600 (Cytiva) with 25 mM Tris pH 7.5 and 150 mM NaCl to obtain diUb A. K48-linked diUb B was generated by incubating 500 μM Ub wt and 500 μM Ub-LAT* (bearing the Sortase 2A motif and a H6 tag at its C-terminus) with 5 μM UBE2R1 and 50 nM UBA1 in 50 mM Tris pH7.5, 10 mM MgCl2-ATP. After overnight incubation at 37 °C the reaction mixture was diluted 1/20 into 20 mM Tris pH 7.5, 300 mM NaCl, 30 mM Imidazole and applied to NiNTA (Cytiva) to remove Ub wt and Ub wt based K48-chains followed by size exclusion chromatography using a Superdex 75 16/600 (Cytiva) with 25 mM Tris pH 7.5 and 150 mM NaCl to obtain diUb B. K48-linked tetraUb was generated by incubating 300 μM diUb A and 200 μM diUb B with 5 μM UBE2R1 and 100 nM UBA1 in 50 mM Tris pH7.5, 10 mM MgCl2-ATP. After incubation for 6 h at 37°C the reaction mixture was diluted 1/20 into 20 mM Tris pH 7.5, 300 mM NaCl, 30 mM Imidazole and applied to NiNTA (Cytiva) to remove diUbA and diUbA based K48-chains followed by size exclusion chromatography using a Superdex 75 16/600 (Cytiva) with 25 mM Tris pH 7.5 150 mM NaCl as buffer to obtain K48-tetraUb bearing the Sortase 2A motif at the C-terminus of the most proximal Ub moiety. Charging of K48-tetraUb onto sfGFP-K215GGK-CycB was carried out by incubating 20 μM sfGFP-K215GGK-CycB with 10 μM K48-tetraUb in presence of 5 μM Sortase 2A. After overnight incubation at 4 °C 200 μM PVS were added to the reaction mixture for an additional hour at 4 °C for Sortase 2A inhibition. Afterwards the reaction mixture was diluted 1/20 into 20 mM Tris pH 7.5, 300 mM NaCl, 30 mM Imidazole and applied to NiNTA (Cytiva) to remove Sortase 2A followed by size exclusion chromatography using a Superdex 75 10/300 (Cytiva) with 25 mM Tris pH 7.5 and 150 mM NaCl to obtain K48-tetraUb-sfGFP-K215GGK-CycB.

### Generation of branched K48-K11 ubiquitin chains on Securin

Securin modification with K48-K11 chains was achieved through APC/C-dependent ubiquitination reactions. Briefly, these recombinant proteins were purified as previously described (*35*). After mixing 1μM UBA1, 5μM UBCH10/UBE2C, 10mM MgCl2-ATP, 1μM CDH1, 100 nM APC/C, 10 μM Securin, the reactions were started by adding 100 μM ubiquitin. After 1.5 hours at room temperature, the reactions were quenched with 50 mM EDTA, aliquoted, and flash frozen. Ubiquitinated securin was purified by size-exclusion chromatography using 50 mM HEPES 7.5, 0.15M NaCl, 0.5 mM TCEP prior to adding it to the proteasome for structural studies.

### Proteasome purification

HEK293 cells expressing hexahistidine, TEV cleavage site, biotinylation site, hexahistidine (HTBH)-tagged PSMD14, a gift from L. Huang (*36*) were dounce-homogenized in lysis buffer (25 mM BisTris pH 6.5, 50 mM KCl, 5 mM MgCl2, 10% glycerol, 5 mM ATP, 0.5 mM TCEP, 0.1 mM PMSF, 1x Benzonase, 1% NP40) containing 1× protease inhibitor cocktail (Roche cOmplete). The lysate was cleared by centrifugation (1 hour, 50,000 x g, 4°C) and filtered using Miracloth and 0.45 μm filters.

Using an ÄKTA pure system (Cytiva), the cleared lysate was loaded onto a 1 mL HP Streptavidin column (Cytiva) equilibrated with 26S proteasome buffer (25 mM BisTris, pH 6.5, 50 mM KCl, 5 mM MgCl2, 10% glycerol, 5 mM ATP, 0.5 mM TCEP). The column was washed with 10 column volumes (CV) of 26S proteasome buffer. Proteasomes were cleaved from the column overnight using GST-tagged TEV protease.

To remove GST-tagged TEV protease, a 5 mL GSTrap HP column (Cytiva) equilibrated with 26S proteasome buffer was used. Cleaved 26S proteasomes were eluted with 26S proteasome buffer, and GST-TEV protease was eluted separately with GST-elution buffer (25 mM BisTris, pH 6.5, 50 mM KCl, 5 mM MgCl2, 10 mM GSH, adjusted to pH 7.0).

Western blotting was performed to detect RPN1 and GST in the 26S proteasome and GST elutions using anti-RPN1 antibody (ab140675, 1:10,000 dilution) and anti-GST antibody (RPN1236, 1:10,000 dilution). 26S proteasome concentrations were quantified using the Bradford assay with a BSA standard curve. Their degradation activity was tested with poly-Ub[K48]-GFP-35 (see below), and their composition was analyzed by tandem MS.

### Grid preparation

Immediately before grid preparation, all 26S proteasome samples were buffer exchanged into 26S proteasome buffer without glycerol in 4 mM ATP or ATPγS using Zeba Spin Desalting Columns (0.5 mL, 7K MWCO, 89882).

On an in-house built manual plunge freezing device at RT (20°C, 40% humidity), 3 μL of 66.6 nM 26S proteasome was pipetted on a holey carbon R3.5/1 Cu200 grid (Quantifoil) covered with a 2 nm continuous carbon foil that was plasma cleaned by a SCD005 sputter coater (Bal-Tec, 60 s, 25 mA). 1 μL of 10 μM Tetra-Ub[K48]-GFP-CycB or Poly-Ub[K11/K48]-Securin was added, mixed by rapid pipetting, followed by immediate manual blotting (Whatman grade 1) and plunge freezing into liquid ethane. The total reaction time from mixing to freezing was measured by an assistant with a stopwatch and was 3 s.

### CryoEM screening and data collection

CryoEM grids were screened on a 200 kV Glacios TEM (Thermo Fisher) at the VBCF EM facility. Grids with good ice quality and intact 26S proteasome particles were selected for data collection on a 300 kV Titan Krios G4 TEM (Thermo Fisher) at the IMP. This instrument was equipped with an E-CFEG electron source, a Selectris X imaging filter (Thermo Fisher) with a 10 eV slit width, and a Falcon 4i direct electron detector. Data collection was performed using EPU (Thermo Fisher) with 40 frames per exposure, a pixel size of 0.951 Å²/px, a total dose of 40 e^−^/Å², and a defocus range of −0.6 to −2 μm in −0.2 μm increments.

### CryoEM image processing

All datasets were pre-processed using CryoSPARC v4.6 (*37*), including Patch Motion Correction, Patch CTF Estimation, Curate Exposures, Particle Template Picking, Particle Extraction, 2D Class Cleaning, Ab-initio Reconstruction, and Volume Alignment Tool. Heterogeneity sorting was performed with CryoSPARC v4.6 (Heterogeneous Refinement and 3D Classification), RELION v4.0 (*38*) (3D Classification), or cryoDRGN v0.3.4 (*39*) (Heterogeneous Reconstruction). Final 3D refinements were conducted using CryoSPARC v4.6 (Homogeneous Refinement and Local Refinement).

### Functional genetic assays

RKO iCas9 cells (*40*) were either engineered to ectopically express mCherry-MYC (*41*) or mCherry-Securin and single cell derived clones were tested for reporter function. Codon-optimized and sgRNA-resistant proteasome subunit cDNAs were synthesized by Twist Biosciences and cloned into a lentiviral overexpression vector (pLV-EF1a-IRES-Puro; Addgene #85132) and packed into virus particles. Transduced reporter cells were fully selected with Puromycin (Sigma-Aldrich). For proliferation experiments, an equal number of cells were sparsely seeded in 24-well plates and harvested 72 hours later. Cell numbers were determined using a Novocyte Penteon flow cytometer (Agilent). For fluorescent reporter assays, fully selected cells were transduced with dsgRNA-Thy1.1-Neo (*41*) targeting the respective proteasome subunit. 2 or 4 days after Cas9 induction, cells were immunostained for Thy1.1 expression and Fluorescence intensity of mCherry quantified in sgRNA expressing cells.

### Generation of K63-, K48-linked or K11/K48 branched ubiquitin chains on Ub-GFP-35

Ub-GFP-35 (a gift from A. Matouschek) was used as model substrate to measure 26S proteasomal degradation rates depending on different ubiquitin modifications. (*13*) Modification with K48 linked chains was achieved by ubiquitination reactions containing 100 μM Ub-GFP-35, 1.5 μM UBA1, 1 μM Super K48 chain former (a gift from C. S. Coleman), 10 mM ATP-MgCl_2_ and 1 mM Ubiquitin in ubiquitination buffer (20 mM HEPES, pH 7.4, 50 mM NaCl) were setup overnight (ON) at 30 °C. Negative control reactions were performed either without Ub-GFP-35, without ATP or with ubiquitin K48R. Species with 1-3, 4-7, 6-10, or 8-11 ubiquitin moieties were purified by size exclusion (Superdex 200 16/600).

Modification with K11/K48 chains was achieved through APC/C-dependent ubiquitination reactions containing 10 μM Ub_6-10_[K48]-GFP-35, 1.5 μM UBA1, 5 μM UBE2S, 1 μM CDH1, 100 nM APC/C, 10 mM MgCl_2_-ATP and 1 mM ubiquitin. Negative control reactions were performed either without ATP or with ubiquitin K11R.

Modification with K63 linked chains was achieved as previously described. (*13*) In brief, ubiquitination reactions containing 100 μM Ub-GFP-35, 1.5 μM UBA1, 1 μM UBC13, 1 μM MMS2, 10 mM ATP-MgCl_2_ and 1 mM Ubiquitin in ubiquitination buffer were setup overnight (ON) at 30 °C. Negative control reactions were performed either without Ub-GFP-35, without ATP or with ubiquitin K63R. Species with 6-10 ubiquitin moieties were purified by size exclusion (Superdex 200 13/300).

### 26S proteasome degradation assays

To measure fluorescence changes over time, the reactions were performed in a microplate fluorescence reader in 26S activity buffer (25 mM Tris, pH 7.5, 4 mM MgCl2, 4 mM ATP, 5% glycerol, 1 mM DTT, 2 g/L BSA). In triplicates, 25 μL of Mix2 (100 nM 26S proteasome), 50 μL blanks (26S activity buffer only), or 50 μL substrate-only (50 nM Ub-GFP-35) were pipetted into a microplate (Greiner, 384 well, PS, F-Bottom, Black, Non-Binding, Item No.: 781900) pre-incubated with 26S activity buffer for 60 minutes at 37°C and dried with compressed air. Using a Synergy H1 (BioTek) microplate fluorescence reader preheated to 37°C, the substrate-only samples were used for automated z-height autofocus and to set the detector gain for 488 nm excitation and 520 nm emission. Reactions were initiated by adding 25 μL of Mix1 (100 nM mono-Ub-GFP-35 or poly-Ub-GFP-35 with K63, K48 or K11/K48 ubiquitin chains) and fluorescence was measured every minute for 3 hours. Linear regression models were fitted to the linear phase of the reaction using Python. Initial degradation rates (V0 [nM/min]) were calculated, and the averages ± SD were plotted.

To monitor the degradation of poly-Ub-GFP-35 with K63, K48 or K11/K48 ubiquitin chains on gel, 4-20% SDS-PAGEs were run an imaged on a ChemiDoc imaging system using fluoresceine read out.

## Acknowledgments

We acknowledge Anton Meinhart, Zuzana Hodakova, Bojan Zagrovic and Jan Michael Peters for helpful discussions on the manuscript.

## Funding

We thank Derek Bolhuis and Liying Cui for their help with generating ubiquitinated securin. KL is grateful for funding by the European Research Council (ERC) under the European Union’s Horizon 2020 research and innovation program (grant agreement no. 101003289—Ubl-tool) as well as funding from ETH Zurich Our work is supported by NIH R35GM128855 and UCRF (NGB); Research in the laboratory of DH is supported by Boehringer Ingelheim, the Austrian Research Promotion Agency (Headquarter grant FFG-852936), the Vienna Science and Technology Fund (grant LS19-029) the Austrian Science Fund (FWF) **10.55776/**project number FWF. For the purpose of Open Access, the author has applied a CC BY public copyright license to any Author Accepted Manuscript (AAM) version arising from this submission.

### Author contributions

Conceptualization: SJA, DH Methodology: SJA, RWK, MF Investigation: SJA, RWK, MF, HK, IG Visualization: SJA, RWK, MF Funding acquisition: JZ, KL, NGB, DH Project administration: DH Supervision: JZ, KL, NGB, DH Writing – original draft: SJA, DH Writing – review & editing: SJA, RWK, MF, HK, KL, NGB, DH

## Data and materials availability

Cryo EM maps have been deposited on the EMDB on the accession numbers XXXX and XXXX. The corresponding pdb files have been deposited at the pdb. Raw micrographs and particles stacks have been deposited to the EMPIAR.

---

**Supplementary Table S1:**
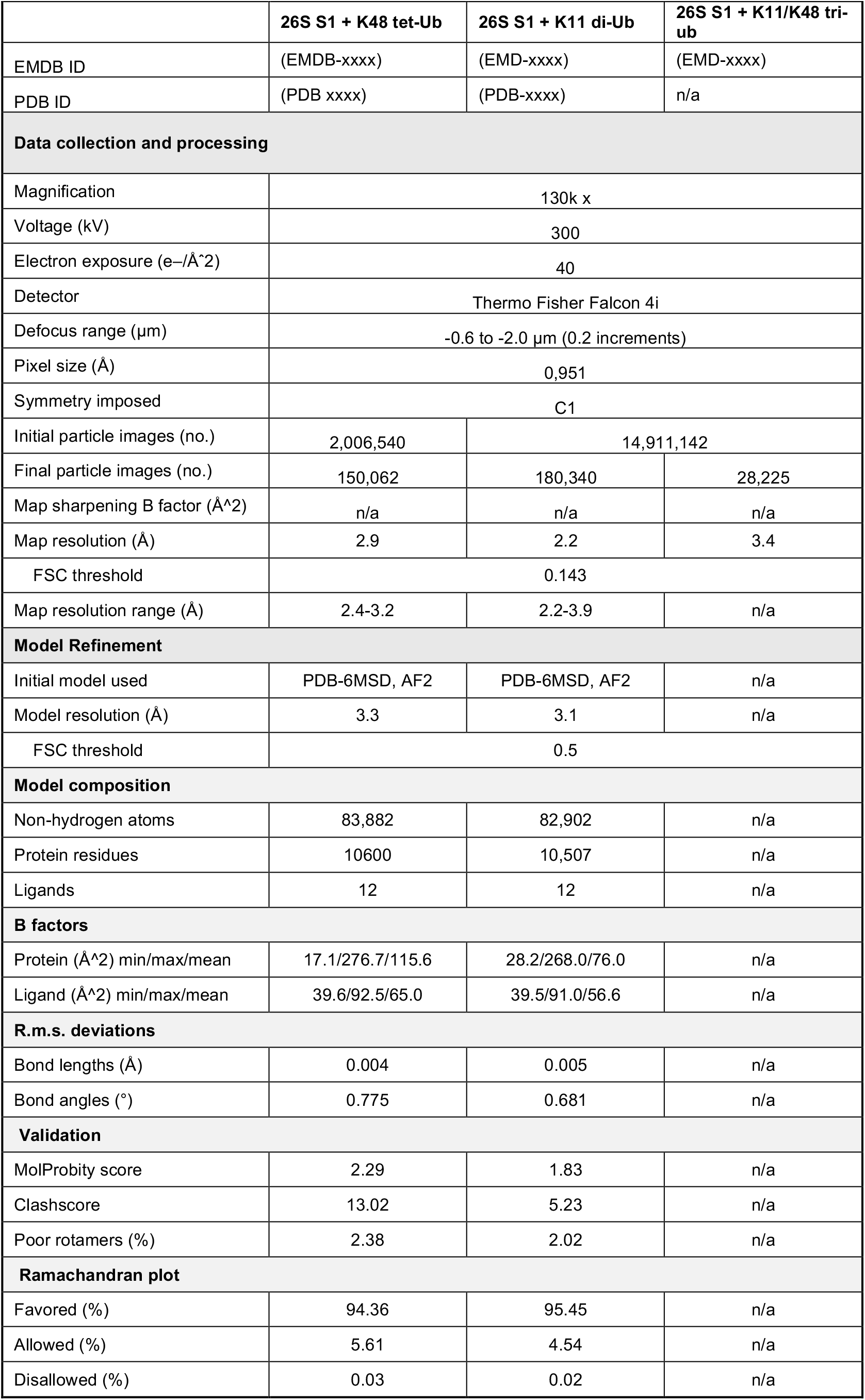
Statistics of cryo-EM data collection, processing and model refinement.

**Supplementary Fig. S1.**
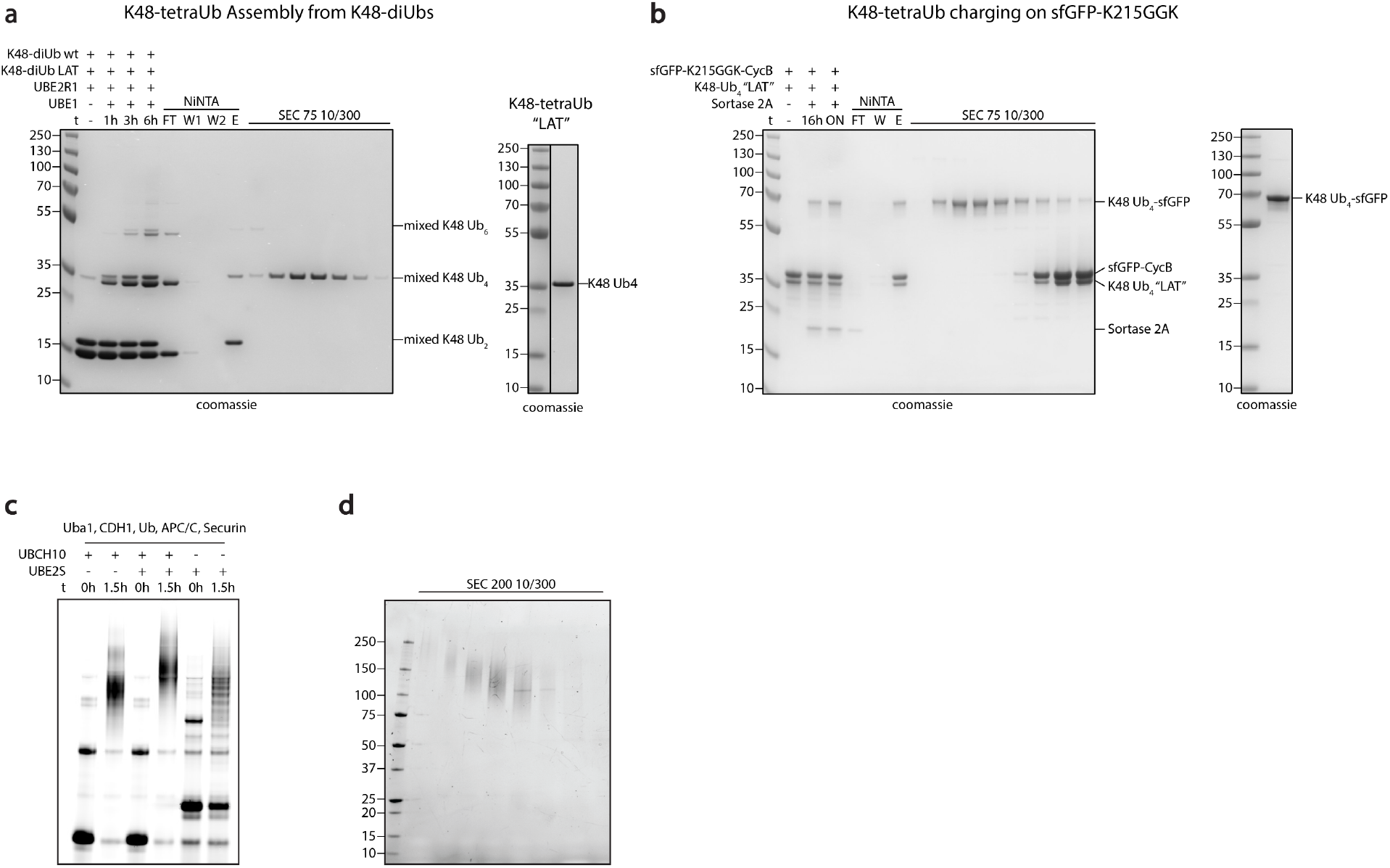
K48 linked tetra-Ub synthesis and K11/K48 branched chain ubiquitination. (a) Left: SDS-PAGE analysis of UBE2R1-mediated K48-tetraUb formation using diUbs A/B. NiNTA chromatography followed by size exclusion chromatography yields K48-tetraUb displaying a Sortase2A motif at the most proximal Ub moiety (FT = flow through, W = wash, E = elution). Right: SDS-PAGE analysis of purified K48-tetraUb. (a) Left: SDS-PAGE analysis showing Sortase2A-catalyzed charging of K48-tetraUb onto the non-canonical amino acid bearing sfGFP-K215GGK-CycB. NiNTA chromatography followed by size exclusion chromatography yields K48-tetraUb-sfGFP-K215GGK-CycB (FT = flow through, W = wash, E = elution). Right: SDS-PAGE analysis of purified K48-tetraUb-sfGFP. (c) APC/C ubiquitination reactions of securin using UCH10 for K48-linked chain formation, UBE2S for K11-linked chain formation, or both forming K11/K48 branched chains. (d) Purification of K11/K48 branched chain ubiquitinated securin by size exclusion.

**Supplementary Fig. S2.**
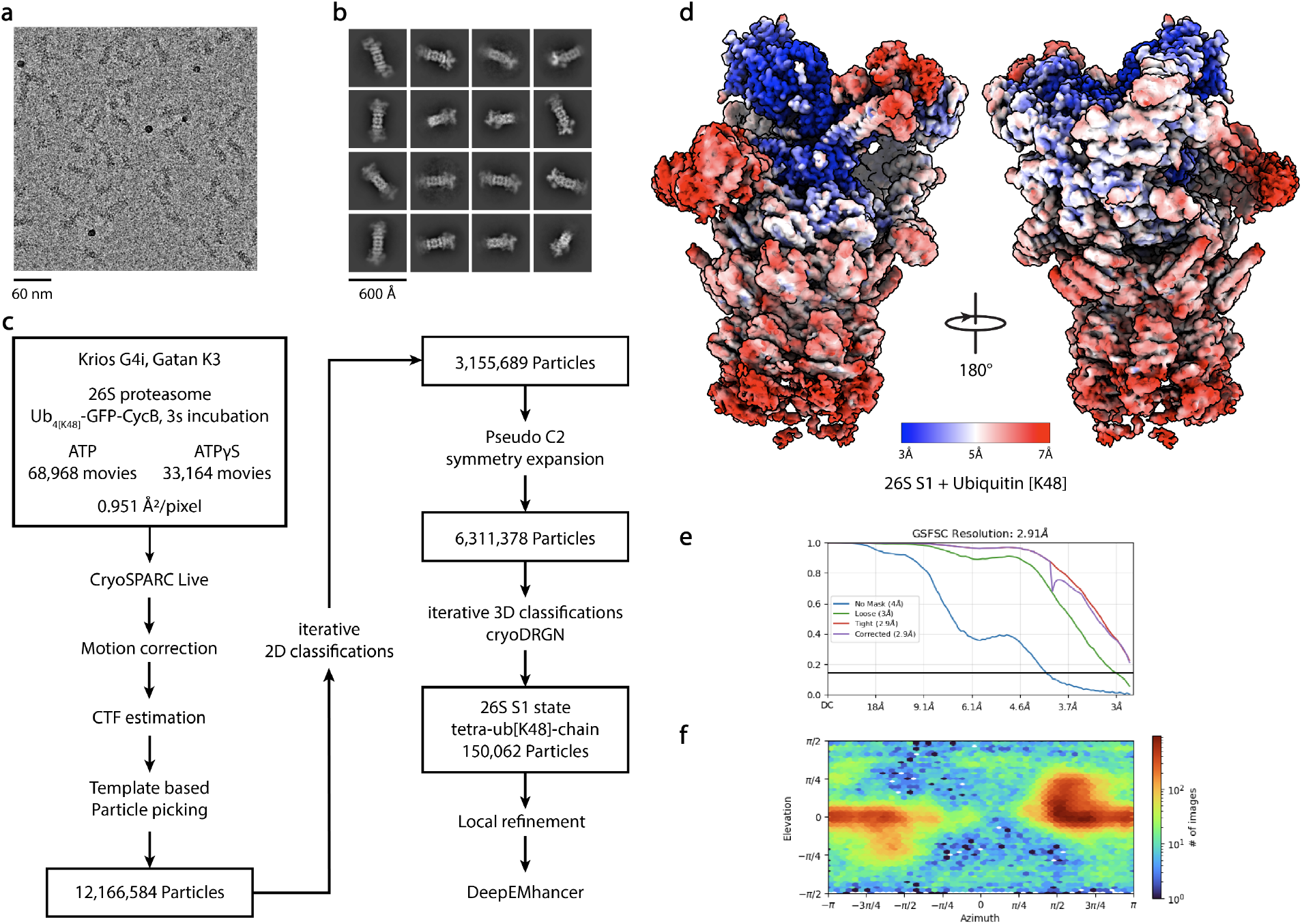
CryoEM analysis of 26S proteasome interacting with K48-linked tetra-Ubiquitin. (a) Representative micrograph. (b) Representative 2D class averages. (c) Processing pipeline. (d) Local resolution estimation of local refined map. (e) GSFSC curve of local refined map. (f) Viewing angle distribution.

**Supplementary Fig. S3.**
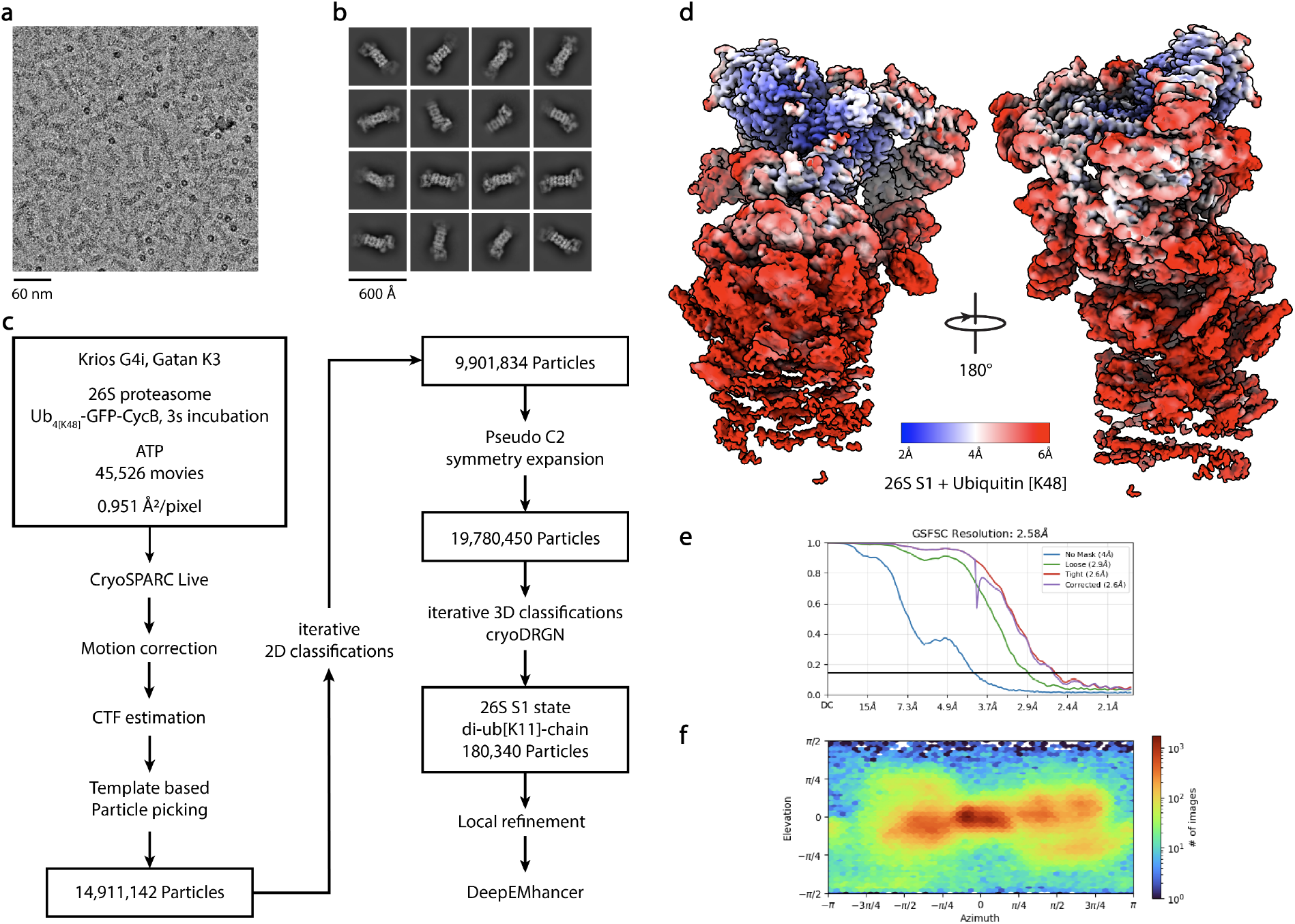
CryoEM analysis of 26S proteasome interacting with K11 linked di-Ubiquitin. (a) Representative micrograph. (b) Representative 2D class averages. (c) Processing pipeline. (d) Local resolution estimation of local refined map. (e) GSFSC curve of local refined map. (f) Viewing angle distribution.

**Supplementary Fig. S4.**
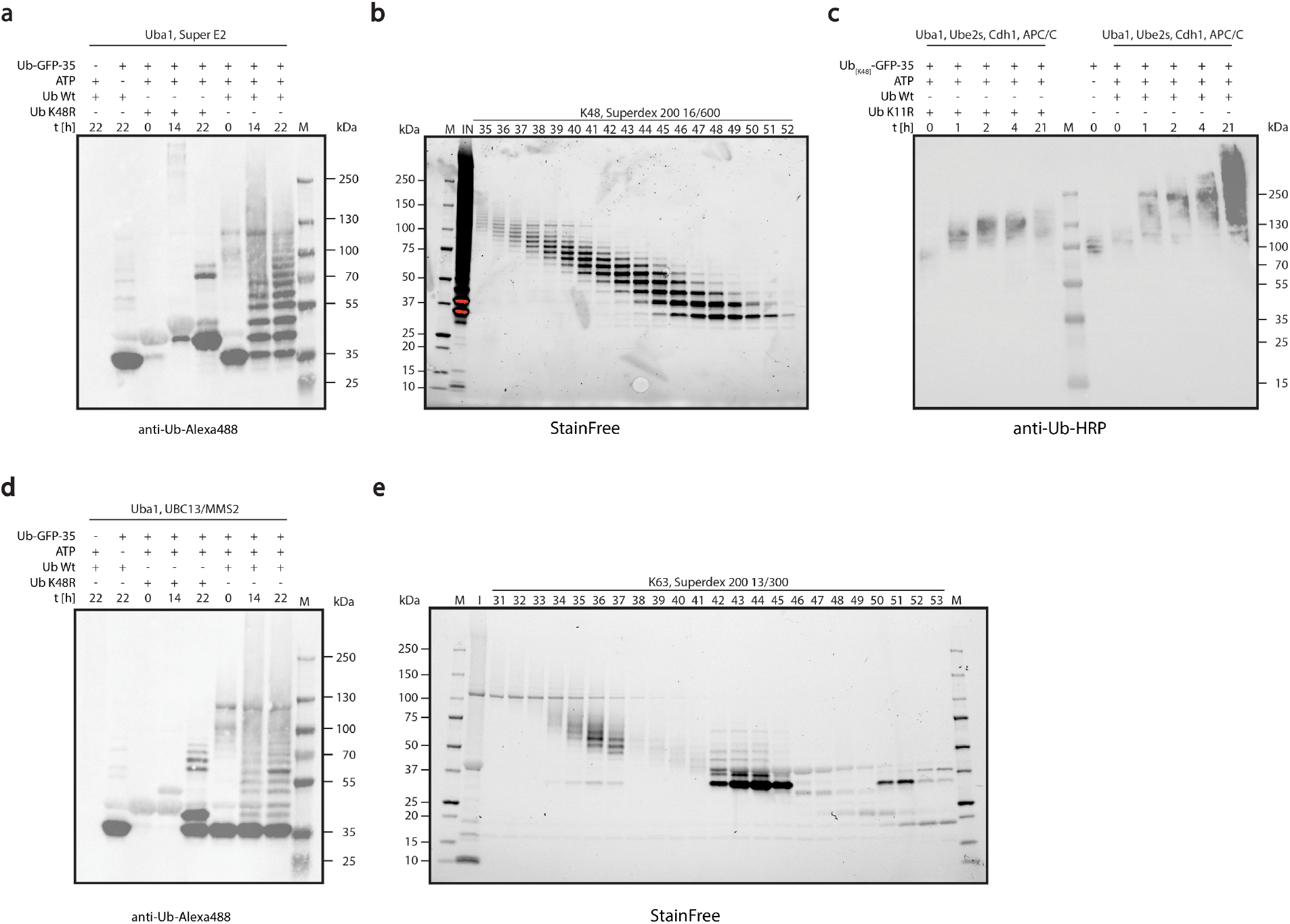
Ubiquitination of Ub-GFP-35 model substrate. (a) Western blot against ubiquitin of ubiquitination reaction of Ub-sfGFP(CP8)-35 with Uba1, Super E2, ubiquitin WT or ubiquitin K48R as control. (b) SDS-PAGE of size exclusion chromatography of ubiquitinated Ubn_[k48]_-sfGFP(CP8)-35 to separate the substrate according to the ubiquitin chain length. (c) Western blot against ubiquitin of ubiquitination reaction of Ub_6-10[K48]_-sfGFP(CP8)-35 with Uba1, UBE2S, APC/C, CDH1, ubiquitin WT or ubiquitin K11R as control. (d) Western blot against ubiquitin of ubiquitination reaction of Ub-sfGFP(CP8)-35 with Uba1, UBC13/MMS2, ubiquitin WT or ubiquitin K63R as control. (e) SDS-PAGE of size exclusion chromatography of ubiquitinated Ub_n[K63]_-sfGFP(CP8)-35 to separate the substrate according to the ubiquitin chain length.

**Supplementary Fig. S5.**
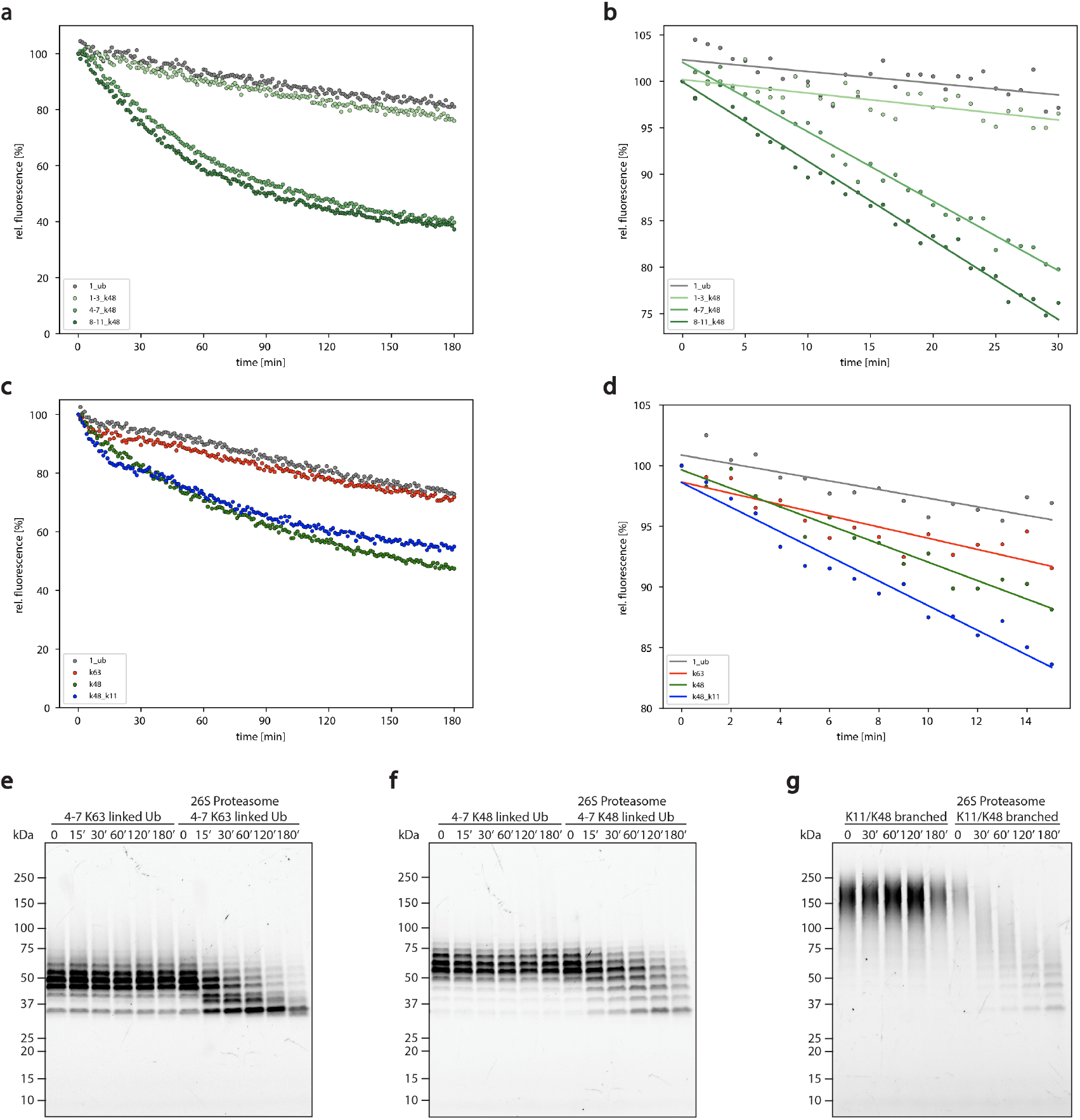
In vitro 26S proteasome degradation assays of Ub-GFP-35 model substrate depending on the Ubiquitin chain type. (a) 26S proteasome degradation activities of Ub-GFP-35 and Ub_[K48]_-sfGFP(CP8)-35 with different chain lengths (n = 3). (b) Linear fits of (a) to determine the initial degradation rates (n = 3). (c)26S proteasome degradation activities of Ub_n_-GFP-35 with different ubiquitin chain types (n = 3). (d) Linear fits of (c) to determine the initial degradation rates (n = 3).(e) 26S proteasome degradation activity of Ub_[K63]_-sfGFP(CP8)-35 with 4-7 ubiquitin moieties, with or without proteasome (f) 26S proteasome degradation activity of Ub_[K63]_-sfGFP(CP8)-35 with 4-7 ubiquitin moieties, with or without proteasome (g) 26S proteasome degradation activity of Ub_[K11/K48]_-sfGFP(CP8)-35, with or without proteasome.

**Movie S1. Engagement of proximal ubiquitin.**

Model morph of the K48-linked tetra-ubiquitin chain in a pre-engagement state interacting with the 26S proteasome (this study) and a proximal ubiquitin relative to the substrate engaged with the 26S proteasomal deubiquitinase RPN11 (6MSD) with superimposed Ub2, Ub3 and Ub4 (this study).

**Movie S2. Simulated conformational changes of the 26S proteasome interacting with a K48-linked tetra-ubiquitin chain**

Model morph of the K48-linked tetra-ubiquitin chain in a pre-engagement state interacting with the 26S proteasome (this study) the 26S proteasome with engaged ubiquitin in the S1 state (6MSD), S2 state (6MSE), S2-S3 intermediate state with RPT4/5 passing by Ub2 (6MSE, 6MSG), S3 state (6MSG) and S3 state after deubiquitination (6MSH).

